# Discovery of 10,828 new putative human immunoglobulin heavy chain IGHV variants

**DOI:** 10.1101/2021.01.15.426262

**Authors:** Fabio R Martins, Lucas Alves de Melo Pontes, Tiago Antônio de Oliveira Mendes, Liza F. Felicori

## Abstract

The correct identification of immunoglobulin alleles in genome sequences is a challenge. Nevertheless, it can assist in the study of several human diseases associated with the antibody repertoire and in the development of new therapies using antibody engineering techniques. The advent of next-generation sequencing of human genomes and antibody repertoires enabled the development of several tools for the mapping and identification of new immunoglobulin (Ig) alleles. Some of these tools use 1,000 Genomes (G1K) data for new Ig alleles discovery. However, genome data from G1K present low coverage and variant call problems. Here, a computational screen of immunoglobulin alleles was carried out in the Genome Aggregation Database (gnomAD), the largest high-quality catalogue of variation from 125,748 exomes and 15,708 human genomes.

A total of 10,909 putative IGHV alleles were identified, in which 10,828 of them are new and 2,024 appear at least in 6 different alleles from genomes/exomes. The IGHV2-70 was the IGHV gene segment with the largest number of variants described. The majority of the variants were found in the framework 3 and most of them are missense. Interestingly, a large number of variants were found to be population exclusive. A database integrated with a web platform was created (YGL-DB) to store and make accessible the likely new variants found.

This available data can help the scientific community to validate new IGHV variants as well as it can shed light on the importance of variants in disease development and immunization protocols.

## Introduction

According to the international ImMunoGeneTics information system (IMGT) the human immunoglobulin heavy chain (IGH) locus at 14q32.33 comprises mostly 38-46 functional IGHV genes belonging to 7 subgroups, 23 IGHD, 6 IGHJ, and 9 IGHC genes (1).

Variations of these individual immunoglobulin germline genes can influence the B cell receptor repertoire, giving rise to differences in immune response, which may impact disease susceptibility or development. For example, previous studies have shown the association of specific germline genotypes with susceptibility to autoimmune diseases such as rheumatoid arthritis (RA), systemic lupus erythematosus (SLE), and multiple sclerosis (MS) (2, 3).

It should be noted that our understanding of sequence variation for IGHV gene alleles is very limited and even nor accurate (4). Variant analysis is hampered due to repetitive nature of locus, the similarity between the genes, the difficulty to distinguish polymorphisms from somatic hypermutations, high fragmentation of genome data available and finally, IGHV sequence diversity have been based on a restricted ethnic heterogeneity (5, 6).

Some methods have been developed over the years with the aim of discovering new IGHV alleles either utilizing Adaptive Immune Receptor Repertoire sequencing (AIRR-seq) or genomic data (7–12). One of the major limitations of all of these methods is the differentiation between somatic hypermutation and new alleles, which requires further validations. In addition, attempts to immunoglobulin new variant discovery from genome studies use 1,000 Genomes (G1K) data, which analyzed 2,504 genomes from 26 populations. However, genomes sequenced in G1K project presented low coverage and variant calls had higher failure rates using alternative variant discovery/genotyping methods (13).

In this this work, we used data from the Genome Aggregation Database (gnomAD), the largest high-quality catalogue of variation in which variant call is performed using a standardized BWA-Picard-GATK pipeline. This database initially aggregated whole exome sequencing data from 199,558 individuals and whole genome sequencing data from 20,314 individuals. However, after stringent sample quality control, the gnomAD v2.1.1 contains data from 125,748 exomes and 15,708 human genomes, superior to G1K that contained only genome data from 2,504 (14).

Using this high-quality dataset, we were able to describe 10,909 putative IGHV variants whose data can be accessed in our database YGL-DB. This database is the most comprehensive collection of putative IGHV variants currently available. This available data can help the scientific community to validate new IGHV variants including the design of new primers to amplify specific or families of variants or even to validate new variants found from AIRR-seq data.

## Results

### 1. Large-scale human IGHV-gene mining revealed 10,824 new putative IGHV variants

Using the pipeline developed in this paper, 40 functional IGHV genes were found in gnomAD GENCODE annotation (26). After filtering for variants present only in V-REGION, a total of 10,909 variants were found (Table 1).

**Table 1:** Number of times which the variants were sequenced in genome/exome (allele count)

|  |  | Presence of variants in other GLDB |  |  | Type of Mutation |  |  |  |  |  |  |
| --- | --- | --- | --- | --- | --- | --- | --- | --- | --- | --- | --- |
| Allele Count | Number of Variants | IMG T | IgPD B | IMGT/IgP DB | Missense | Synonymous | Stop_gained | Frameshift | Deletion | Insertion | Protein_altering |
| 1 | 4,993 | 4 | 4 | 0 | 3,365 | 1,178 | 172 | 209 | 36 | 27 | 6 |
| 2 | 1,961 | 0 | 2 | 0 | 1,349 | 522 | 53 | 30 | 3 | 3 | 1 |
| 3 | 974 | 1 | 0 | 0 | 678 | 262 | 18 | 11 | 4 | 1 | 0 |
| 4 | 571 | 1 | 0 | 1 | 379 | 172 | 11 | 8 | 1 | 0 | 0 |
| 5 | 386 | 1 | 1 | 0 | 244 | 124 | 11 | 5 | 0 | 2 | 0 |
| 6-10 | 765 | 2 | 0 | 0 | 514 | 224 | 14 | 11 | 0 | 2 | 0 |
| 11-20 | 417 | 1 | 2 | 0 | 268 | 134 | 7 | 7 | 0 | 1 | 0 |
| 21-30 | 146 | 0 | 0 | 0 | 97 | 41 | 5 | 2 | 1 | 0 | 0 |
| 31-40 | 76 | 0 | 0 | 0 | 49 | 22 | 1 | 4 | 0 | 0 | 0 |
| 41-50 | 42 | 0 | 0 | 0 | 21 | 18 | 1 | 2 | 0 | 0 | 0 |
| 51-60 | 27 | 0 | 0 | 0 | 17 | 8 | 0 | 0 | 2 | 0 | 0 |
| 61-70 | 27 | 1 | 1 | 0 | 12 | 11 | 3 | 1 | 0 | 0 | 0 |
| 71-80 | 16 | 0 | 0 | 0 | 10 | 5 | 1 | 0 | 0 | 0 | 0 |
| 81-90 | 13 | 0 | 0 | 0 | 6 | 7 | 0 | 0 | 0 | 0 | 0 |
| >90 | 495 | 31 | 21 | 11 | 291 | 174 | 8 | 13 | 3 | 6 | 0 |
| Total | 10,909 | 42 | 31 | 12 | 7,300 | 2,902 | 305 | 303 | 50 | 42 | 7 |

Some of the alleles described here were also found in other immunoglobulin germline databases, such as 42 in IMGT/GENE-DB (16), 31 in IgPdb (20) and 12 in both databases. However, none of the variants described here were found on OGRDB (21) and most of them, 10,824, are not present in these current and analyzed GLDBs.

There is a large variability in the number of variants from different IGHV genes ranging from 103 variants found for IGHV1-45 to 807 for IGHV2-70 (Supp Figure 1). Most of the variants, 4,634, were found in the largest IGHV subgroup, IGHV3 (Supp Table 1).

At least one variant from each V gene was already described in other germline databases (GLDB) such as IMGT and IgPDB (except variants from IGHV1-3, IGHV3-43 and IGHV3-66). However, most of the putative variants are new (Table 1, Supp Figure 1). Variants were also found distributed all along IGHV locus. Interesting to note that 45.7% (4,993) of the new putative variants described here appear only once in genome/exomes present in gnomAD but some of them were also detected on IMGT or IgPDB (Table 1). Importantly, 2,024 new putative variants appear at least in 6 different alleles (Table 1).

### 2. The majority of the IGHV variants are in Framework 3 region

Interesting to note that the majority (84,3%) of the variants are present in the framework regions: 20.9% or 2280 variants in FWR1, 22% or 2403 in the FWR2 and the large majority in FWR3 (4508 variants or 41.3%). Some of the variants were also found in the regions corresponding to CDR1 (743 variants, 6.8 %), CDR2 (449 variants, 4.1%) or the beginning of CDR3 (526 variants, 4.8%). It’s also interesting to observe that the majority of the variants don’t have a preference for a specific position in V-REGION (Figure 2). The frequency of 1 variant (35%), 2 variants (33%) or even 3 (23%) variants at the same position is elevated. However, few variants, as the case of the variants derived from IGHV4-4 gene present 6 variants at the same position in the FWR3 (Figure 2).

### 3. Most of the IGHV variants described are missense

From 10.909 variants described with this approach, the majority are missense (7,300 variants: 66.9%) and synonymous (2,902: 26.6%) single nucleotide variants (SNVs) (Table 1). IGHV2-70, IGHV1-2 and IGHV1-69 are the IGHV genes involved in the majority of missense and synonymous variations (Supp Figure 2).

One can observe that 305 putative variants found in this work result in stop codon (stop_gained). Including 8 of them present in more than 90 alleles. In addition, more than 20 frameshifted and in frame deletion or insertion appeared in more than 90 alleles (Table 1). The biggest deletions observed were for variants derived from IGHV6-1 (104 nucleotides), IGHV2-70 (89 nucleotides) and IGHV1-3 (84 nucleotides), all of them in the FWR2. On the other hand, the biggest insertions were observed in the variants derived from IGHV1-18 (with an insertion of 99 and 42 nucleotides) and from IGHV2-5 (84 nucleotides) (Supp Figure 3).

### 4. Large number of variants are unique in different world populations

It was observed that 277 variants were shared by all the populations present in gnomAD (African, Latino, East Asian, South Asian, European (Finish and non-Finish) and Ashkenazi Jewish). Most of these variants are highly prevalent, as can be seen in Table 2 showing the most prevalent ones.

**Table 2:** Top 10 most prevalent IGHV variants

| IGHV gene name | Variant Description | Allele Count | Allele frequency |
| --- | --- | --- | --- |
| IGHV2-70 | 14-107178832-A-G | 272,720 | 0.9993 |
| IGHV4-31 | 14-106805440-A-G | 270,476 | 0.9999 |
| IGHV4-61 | 14-107095326-C-T | 204,037 | 0.8493 |
| IGHV2-5 | 14-106494269-T-C | 193,598 | 0.6999 |
| IGHV1-3 | 14-106471388-C-A | 173,235 | 0.6265 |
| IGHV1-3 | 14-106471353-C-T | 171,661 | 0.6204 |
| IGHV1-3 | 14-106471268-A-G | 171,004 | 0.6199 |
| IGHV1-3 | 14-106471273-C-T | 167,497 | 0.6099 |
| IGHV2-70 | 14-107178965-A-C | 159,909 | 0.5804 |
| IGHV1-3 | 14-106471528-C-A | 156,122 | 0.5837 |

Interesting to note that more than one highly prevalent variant comes from IGHV2-70 and IGHV1-3 genes. In addition, almost all sequenced individuals have the top 2 variants resulted from a substitution of an adenine to guanine.

One could observe also in this work a high number of unique population variants, but present in low frequency (Table 3). We noticed the presence of 729 exclusive IGHV variants in Africans, 809 in Americans (Latinos), 857 in Asians (476 South Asia + 381 East Asia) and in 3,954 Europeans (3,415 non-Finnishi, 468 Finnishi and 71 Ashkenazi Jewish). The majority of exclusive variants (3,415) were found in Europeans (European non-Finish). However, the most frequent variants of this population are derived from IGHV3-21 gene with frequency of 0,0089% in the populations studied (Table 3). The most frequent unique variant was found in Africans and is derived from IGHV1-24 present in 102 alleles sequenced in gnomAD (frequency of 0.00367%).

**Table 3:** Number of unique variants per population and the most frequent variant of the population

| Population | Number of unique variants | Most frequent IGHV | Variant Description | Allele Count | Allele Frequency |
| --- | --- | --- | --- | --- | --- |
| African | 729 | IGHV1-24 | 14-106733417-A-G | 102 | 0.0003674 |
| Latino | 809 | IGHV1-2 | 14-106452903-G-C | 61 | 0.0002478 |
| EastAsian | 381 | IGHV3-9 | 14-106552504-T-G | 65 | 0.0002846 |
| South Asian | 476 | IGHV3-53 | 14-107048714-G-A | 39 | 0.0001582 |
| European (non-Finnish) | 3,415 | IGHV3-21 | 14-106691736-G-A | 25 | 0.00008996 |
| Ashkenazi Jewish | 468 | IGHV1-2 | 14-106452694-T-C | 16 | 0.00005776 |
| European (Finnish) | 71 | IGHV3-13 | 14-106586381-C-A | 10 | 0.00004069 |
| Total | 6,349 |  |  |  |  |

## Discussion

In order to better understand the allelic diversity of human IGHV genes, we mined the gnomAD variant catalog, and as a result, we identified 10,909 immunoglobulin heavy chain V-gene variants (IGHV), being mostly of them potentially functional, which represent a substantial increase in the number of known alleles. All alleles found so far are stored on the YGL-DB web platform.

Some authors claim that the reference (IMGT) for immunoglobulins and TCR is incomplete and has inconsistencies (5, 7). One of the facts that support the incompleteness is that the diversity of the IGHV sequences was based on a restricted set of ethnic groups, and therefore, the data may represent only a part of the allelic diversity in humans (2). As shown in Table 3, our work used gnomAD data from 7 ethnic groups (African, East Asian, Ashkenazi Jewish, European - Non Finnish, European - Finnish, South Asian, Latino, as well as individuals whose ethnic origin was not reported), which can serve as a groundwork for future population-level studies of IGHV diversity. Interesting to note that the majority of alleles described in this work is unique to different populations, including understudied Africans, Latinos and Asians populations.

In addition, the germline database expansion produced by our work can benefited IGHV gene analysis software, overcoming problems related to the incompleteness of the reference database. Firstly, it might minimize the erroneous assignment of germline polymorphisms as somatic hypermutations (SHMs) by repertoire annotation pipelines Secondly, this expanded database could be used as a training set for statistical tools (such as profile HMMs) to identify new functional V genes, pseudogenes and orphans in genome sequences.

Moreover, since many IGHV genes and alleles are knowable associated with the susceptibility to diseases, such as: multiple sclerosis (IGHV2), rheumatoid arthritis (IGHV3-30, IGHV4-31, IGHV1-69), systemic lupus erythematosus (IGHV3-30, IGHV4-31), type 1 diabetes (IGHV2, IGHV4, IGHV5), and Kawasaki disease (IGHV1-69, IGHV2-70) (as reviewed by (3)) the analysis of the newfound variants associated with healthy status of individuals can be fruitful.

Furthermore, the increase in the number of known alleles from each IGHV gene allows the identification of comprehensive consensus sequences between different alleles. This is especially useful for the design of multiplex PCR primers for repertoire studies, improving the probability that a primer is capable of amplifying an immunoglobulin transcript sequence of an individual, independently of the previous knowledge of its germline gene repertoire.

It is important to note that the scope of the variant dataset provide by this work is limited by some restrictions: first, the GENCODE GRCh37 reference genome includes only a part of the known IGHV genes (40 of 55) and families (6 of 7 - see Supp Table 1); second, we do not accessed directly the individuals’ genomic information that make up the gnomAD database, which impedes the genotyping of the individual’s alleles; third, the present analysis was restricted to the V-REGION sequences, which narrows the allele’s functionality analysis to checking for the presence of stop codons before the last codon; fourth, we do not know the cell types utilized for the whole genome and exome sequencing for each individual in the dataset. However, besides the limitations, this work describes the most comprehensive collection of putative IGHV variants currently available what can help the scientific community to validate new IGHV variants found from genomic studies and from AIRR-seq data.

## Methods

### Development of a pipeline for IGHV variants discovery from genome and exome

We aimed to identify possible variable (V) gene variants from human heavy chain immunoglobulin (IGHV) germline gene using the gnomAD (14). Since gnomAD (14) adopts the GENCODE release 19 for gene annotation (15) and the standard Immunoglobulin annotation uses the IMGT/GENE-DB (16) as database for germline sequences, a comparison between both were done to extract the correct corresponding position between both datasets.

Briefly, a search in “.gff” file for IGHV positions was done for both NCBI (RefSeq Reference Genome Annotation from build GRCh37 – ref. 17 that uses IMGT standards) and GENCODE (Comprehensive gene annotation from build GRCh37.p13 – ref. 15). Since differences were observed between the positions of each V gene in both databases, all IGHV nucleotide sequences from both dataset were recovered from the related “.fa” files using Samtools v1.6 (http://www.htslib.org/). The sequences for all IGHV from both datasets were aligned using Needleman-Wunsch algorithm through Biopython v1.74 (18). Only the V-REGION (encoded by a V gene: without leader and RSS) was filtered and analyzed. Variant data (sequence and position) were obtained from gnomAD using Selenium library, a web scraping application developed in Java 1.8. For the generation of V-gene new allele, a Java 1.8 program was implemented in which the nucleotide variant recovered from gnomAD v2.1.1 was replaced in the corresponding V-REGION. After nucleotide deletion, addition or substitution in V-REGION, the sequences were complementarily reversed, as standardized by IMGT/V-Quest (19). This methodology is summarized in Figure 1A.

**Figure 1:**
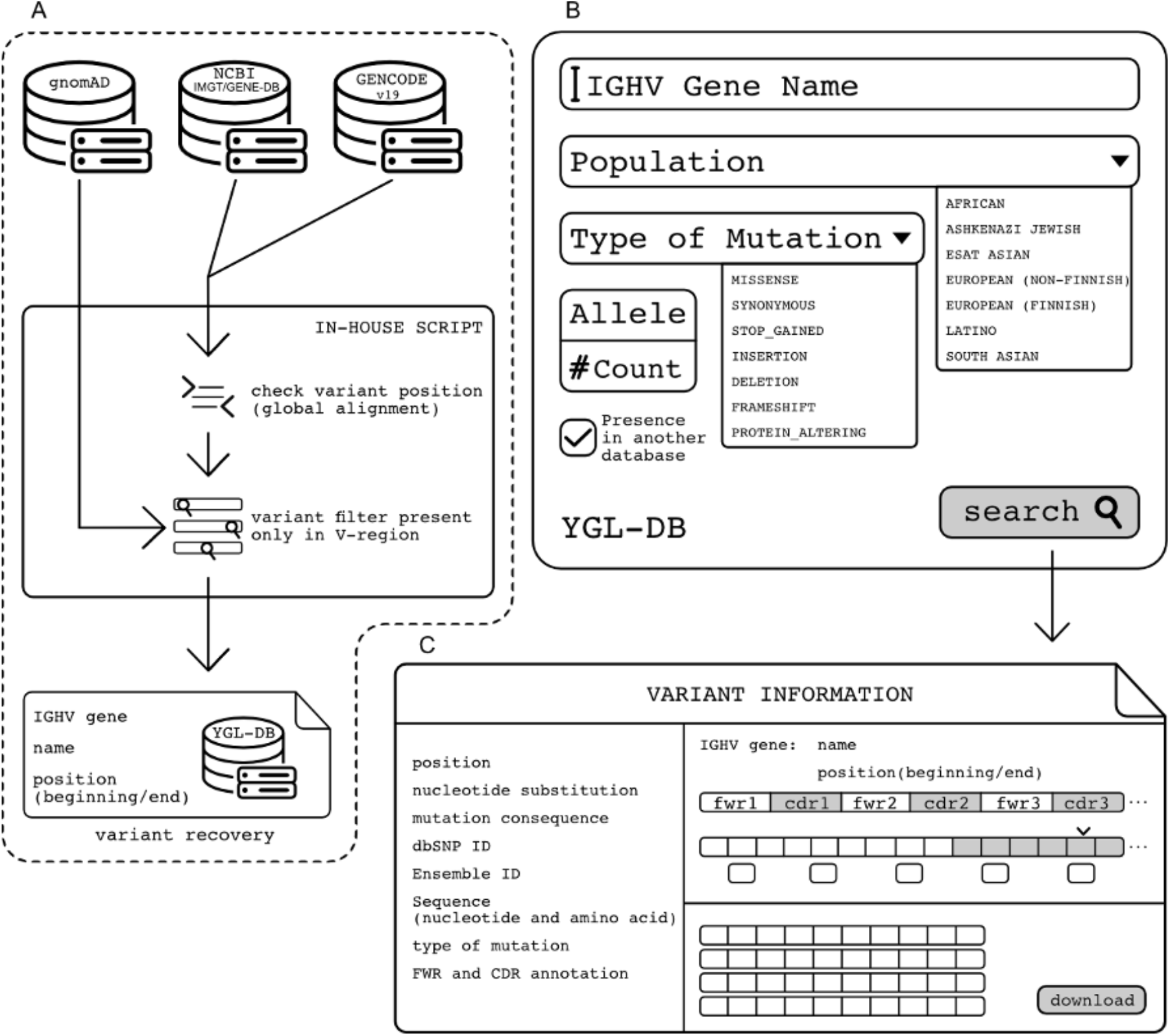
IGHV germline variants discovery strategy and YGL-DB. A) To obtain IGHV variants, IGHV genes names, sequences and positions were extracted from NCBI (that uses IMGT-Gene-DB) and from GENCODE (used by gnomAD). The sequences obtained were compared using global alignment. After manual checking, all the 40 IGHV on gnomAD, which were exactly matched with IMGT sequence were filtered only by it’s V-REGION and the variants were recovered by replacing the corresponding mutant nucleotide in V-REGION. B) YGL-DB input where the user can search the database by IGHV gene name, population, type of mutation, the number of times the variants were sequenced or if the variants are present in other databases. C) YGL-DB output with the corresponding IGHV gene name and position and several variant information: position, nucleotide substitution and consequence in the protein, type of mutation, variant sequence, dbSNP and Ensembl identifier, FWR and CDR annotation in the variants and allele count (number of times this variant were sequenced).

**Figure 2.**
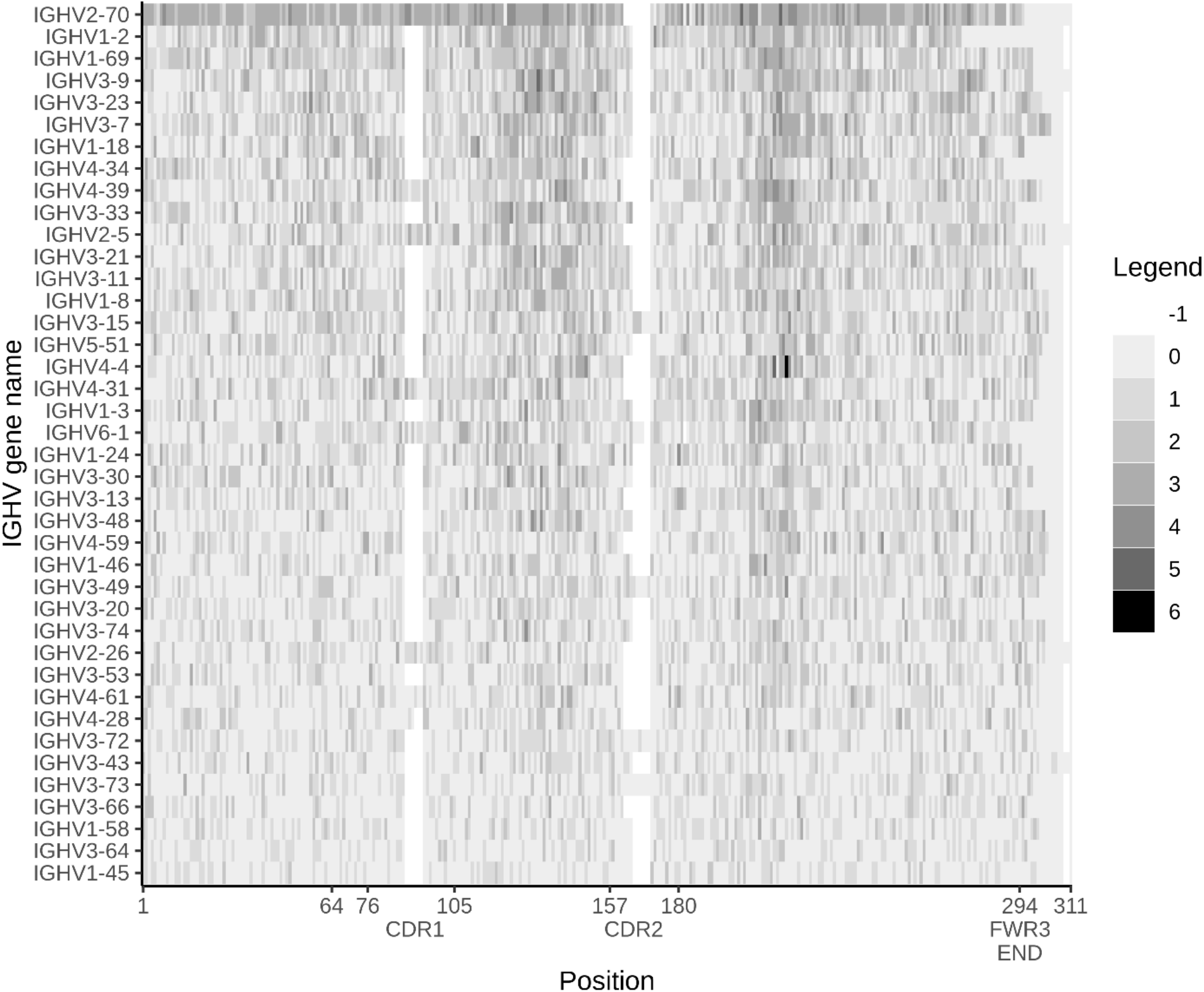
Number of variants in each of the IGHV gene per Position. Heatmap presenting the number of each variant per IGHV position. 0 means no variant in a given position, 1 one variant, 2 for two variants, 3 for three variants, 4 for four variants, 5 for five variants and 6 for six variants in a given position. This heatmap is organized in a decrescent order since the IGHV gene containing the largest number of variants is on top of the figure (IGHV2-70) and the one with fewer variants in the bottom (IGHV1-45).

To better analyze the variants discovered here, all of them were aligned with the corresponding germline IGHV gene using MUSCLE (25) and checked for conserved positions using an in-house Java script.

Each of the new putative alleles found in this work was compared with pre-existed documented alleles from other databases such as IMGT (16), IgPDB (20) and ORGBD (21).

The variants presented in our data and in one or more of the above-described databases were identified using a global alignment (Java 1.8) with sequence coverage and identity of 100%.

## Database Generation

A database named YGL-DB was created containing IGHV gene identification and sequence, position and variant information including: nucleotide sequence, amino acid sequence (obtained through IgBLAST (22)) and position, FWR and CDR position, type of mutation (missense, synonymous, frameshift, in frame deletion or insertion, protein_altering and stop_gained), number of times the variants appear on genome and exomes sequenced in gnomAD, dbSNP and Ensemble IDs (23, 24), the presence of the variant in other databases and the presence in different populations (Figure 1B e 1C).

The database YGL-DB was created using MySQL version 8.0.20 (https://www.mysql.com/) and a Java 1.8 program, in a Linux system (x86_64). For table modeling, MySQL Workbench 8.0.20 was used. YGL-DB is available on: http://bioinfo.icb.ufmg.br/ygl/

## Acknowledgments

Lucas Silva for Figure 1 and YGL-DB design. SynBiom group for fruitful discussion.

